# Grey Mouse Lemurs’ response to acute caloric restriction before reproduction supports the “Thrifty Female Hypothesis”

**DOI:** 10.1101/2023.09.08.556820

**Authors:** Aude Noiret, Fabienne Aujard, Jeremy Terrien

**Affiliations:** Unité Mécanismes Adaptatifs et Evolution (MECADEV), Muséum National d’Histoire Naturelle, CNRS UMR 7179, Brunoy, France

## Abstract

The thrifty female hypothesis states that females would preserve more of their energy reserves during winter than males, because of a sex specific time-frame of energy allocation to reproduction. As males reactivate their reproductive axis before the mating period, while females mainly allocate energy in gestation and lactation, we make the hypothesis that males would have to use shorter torpor bouts and longer periods of normothermic activity in order to promote spermatogenesis during winter, a period of low food availability. Here we applied an acute two-week 80%CR in males and females grey mouse lemurs, shortly before the mating period. We show evidence of thriftier phenotypes in wintering females, which performed deeper and longer torpor bouts than males, and ultimately lost less body mass. Our results thus support the thrifty female hypothesis in a seasonally breeding primate, and reinforce the concept of a sex-biased trade-off in using torpor, which might ultimately benefit to reproduction and survival.

## 1. Introduction

Reproduction and survival are common examples used to depict the concept of trade-off: by investing energy in reproduction, one individual ultimately impacts its own survival chances (Doughty and Shine, 1997; Blomquist, 2009; Cox et al., 2010). In the majority of birds and mammals, the succession of seasons has shaped the necessity to produce offspring in a limited timing and during the most energetically favourable period, i.e. during summer or the wet season (Conover, 1992; Fournier et al., 1999; Ball and Ketterson, 2008). As reproduction is one of the most energy consuming physiological process (Clutton-Brock et al., 1989; Speakman, 2008), some species developed energy-saving mechanisms. Temporal heterothermy is primarily seen as a phenomenon preventing decreased survival chances during winter, when the food availability is low and climate conditions are challenging (Malan and Canguilhem, 1989; Geiser, 2004; Jastroch et al., 2016). By decreasing in a controlled manner their metabolic rate and body temperature -a process called torpor-some organisms manage to prevent excessive fat loss during winter (Geiser, 1998; Melvin and Andrews, 2009). The idea that heterothermy might be used to maximize reproductive success in the long-term by saving energy to allocate later to reproduction is much less described in the literature than the effects of energy reserves allocation to survival. However, torpor has direct costs that could impact short-term survival, as any other stress response (Burton and Reichman, 1999; Prendergast et al., 2002; Luis and Hudson, 2006; Bieber et al., 2014; Wei et al., 2018): in this context, the “torpor optimization hypothesis” states that it should be used only if fat stores are necessary for survival or reproduction (Humphries et al., 2003). In parallel, males and females do not share the same physiological reproductive tasks, when male produce high stocks of spermatozoa before mating, while females ovulate or engage themselves in gestation and lactation. In consequence, female mammals’ reproduction is seen as more pressuring to select for energy saving mechanisms, either because of the high energy expenses allocated to lactation (Clutton-Brock et al., 1989), or because of their more constraining reproductive frame, which consists of investing in their own reserves to achieve reproductive success, in opposition to males’ gamete dispersion in several females (Jonasson and Willis, 2011). Moreover, as survival chances of animals with fast life histories are uncertain, reproductive opportunities are sparce and must be optimized (Sæther et al., 2004), especially in unpredictable environments submitted to high environmental constraint (Dewar and Richard, 2007).

Because of the costs of torpor and the alleged sex-imbalance in energy allocation to reproduction, the “thrifty female hypothesis” (TFH) has been pronounced by Jonasson and Willis (2011). In the little brown bat (*Myotis lucifurus*), the team showed that females retained more fat going through winter than males, which would ultimately confer an advantage in resisting White Nose Syndrome (Jonasson and Willis, 2011). If a link exists between seasonal reproduction and the occurrence of thrifty phenotypes in females, observations of sex differences in energy balance should be made in other heterothermic species. In fact, in polar bears (*Ursus maritimus*), only females can enter hibernation when they are in gestation, depending on environmental conditions (Lennox and Goodship, 2008). Moreover, adult male ground squirrels express shorter torpor bouts than females (Kart Gür and Gür, 2015), but the outcome in body mass is not mentioned. Here, we will focus on confirming the TFH in *Microcebus murinus*, a small nocturnal primate from Madagascar, and seasonal breeder that is well described to use torpor (Giroud et al., 2008; Vuarin et al., 2015).

Evidence for another case of a thriftier female has been shown in this species, as males and females express strong differences in their body mass fluctuations over winter. Torpor event of several days have been observed in wild females, which were rather “fat”, but not in males (Schmid and Ganzhorn, 2009). Indeed, when males begin to lose weight around the middle of the season concomitantly to testicular growth initiation (Perret and Aujard, 2001; Terrien, 2018), females keep their fat storage until summer when mating takes place. Moreover, females were described to use deeper and longer torpor bouts than males to face food rarefaction at the onset of winter in field conditions (Vuarin et al., 2015). However, in a previous study in laboratory conditions, even if wintering males and females expressed strong specificities in their thermic profiles, we have not evidenced a difference in energy balance between sexes under either control ration or a 40% caloric restriction (Noiret et al., 2021). We make the hypothesis that the expression of female thrifty phenotype during winter could occur in more energetically-challenging conditions. Here, we submitted the animals to an intense two-weeks 80% caloric restriction during the second half of winter, when males are known to reactivate spermatogenesis (Perret and Aujard, 2001), and thus placed them in an abrupt change of calorie intake. Indeed, caloric restriction initiating and modulating torpor is widely documented (Giroud et al., 2008; Vuarin et al., 2015). We equipped the mouse lemurs with temperature and activity loggers, monitored their body mass and measured two markers of fitness (cortisol and oxidative DNA damage 8-OHdG) and reproductive success. By putting the animals in a such drastic - yet safe-caloric challenge, we expected to trigger sex-specific energy saving mechanisms, enough to observe differences in body mass loss between males and females *Microcebus murinus*.

## 2. Materials and methods

### Animals and ethical concerns

Twenty-two Grey mouse lemurs (*M. murinus*), 11 males and 11 females all aged from 2 to 4 years and raised in good health in the breeding colony of Brunoy (MNHN, France, license approval n◦ E91-114-1), were included in the experiment. These animals were tested at the end of the winter-like season, during weeks 17 to 21 of winter-like photoperiodic exposure (SD; 10 h light/day), at the time when testis recrudescence is occurring (Perret and Aujard, 2001). Animals were kept in individual cages in semi-isolation from the others (visual, hearing and odorant interactions remaining possible between individuals) in climatic chambers for the duration of the experiment (1 month). Temperature and humidity were maintained constant (24–26◦C and 55%, respectively). The lemurs were fed with a fresh mixture (egg, concentrate milk, cereals, spicy bread, cream cheese, and water), banana, and were provided with ad libitum water. All described experimental procedures were approved by the Animal Welfare board of the UMR 7179, the Cuvier Ethics Committee for the Care and Use of Experimental Animals of the Muséum National d’Histoire Naturelle and authorized by the Ministère de l’Enseignement Supérieur, de la Recherche et de l’Innovation (n◦14075-2018031509218574) and complied with the European ethic regulations for the use of animals in biomedical research.

### Experimental design

We performed an integrative description of physiological parameters, to decipher the energy balance of the animals, their metabolic activity, and their oxidative status. For this, animals were fed daily with a control ration (Control treatment “CTL”) of 22 g of mixture and 3 g of banana (24.48 kcal.day− 1) for 1 week and measured during four consecutive days for indirect calorimetry (Oxymax, Columbus Instrument Inc., Columbus, Ohio, USA). At the end of this procedure, animals’ urine was sampled, always performed ∼3 h prior to lights off. Urine samples were used to measure creatinine, cortisol, and 8-OHdG. After 1 day of recovery, animals were either maintained under CTL diet for 2 more weeks, or fed daily with a 80% reduced ration compared to the CTL condition (Caloric Restriction “CR,” 4.90 kcal.day− 1). Such condition was designed to induce a sufficient caloric stress to challenge the animals, but not enough to put animals at risk considering that animals’ body mass is high at this time of the year. They were monitored with weighing three times a week to follow their body mass (“BM” in g), and measured one time from the tip of the nose to the anus to establish body length (in cm). Individual reproductive state was monitored from morphological changes as previously described (Perret and Aujard, 2001). Males’ testis size was rated at the beginning and the end of the experiment, based on a scale from 0 to 2 depending on testis size and consistency: 0= testes are in the abdominal cavity and scrotal sacs are loose; 0.5 = testes descend and measure 1 cm put together; 1= 2 cm; 1.5 = 3 cm; 2= 4 cm and consistency becomes harder. At the end of the 2-week treatment, they underwent the same procedures as the control period (4-days indirect calorimetry, urine sampling). Animals returned under control diet for several days to allow a full recovery from the CR treatment and returned to their original housing conditions in social groups. Females’ reproductive status was monitored until oestrus by direct visual evaluation, to establish the number of days to mating readiness after photo transition to long days.

### Assessment of metabolic activity and stress response

#### Indirect Calorimetry

We used the same method as described previously (Noiret et al., 2020). Briefly, animals were put in monitored cages for 4 consecutive days to measure their metabolic rate (“MR” in mlO_2_.kg^-1^.h^− 1^, see equation 1, (Lighton, 2018), defined by the oxygen consumption rate (Geiser 2014). We used an automated calorimetric set-up (Oxymax, Colombus Instruments Inc., Columbus, Ohio, USA).

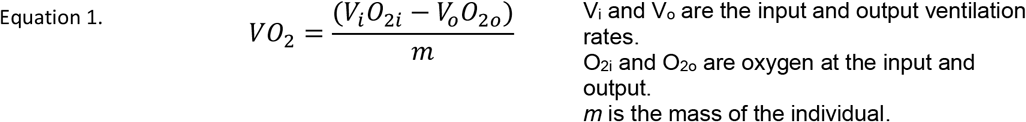

Air was dried with magnesium perchlorate columns prior to analysis. As metabolic rate (“MR”) is acknowledged to be directly linked to oxygen consumption rate (Geiser et al., 2014) we used VO_2_ to describe variations in mouse lemurs’ metabolic rate. Analyses were based on mean parameters over day (period under artificial light) and night (period without artificial light).

#### Body temperature and activity patterns

Mouse lemurs were implanted with DSI telemetry captors (TA-F10, Data Sciences instruments) into their abdominal cavity, to monitor both body temperature (°C) and activity (counts/min). Surgical procedures were performed under general anesthesia (Valium 0.5mg/ 100g; Ketamine 2 mg/ 100g; isoflurane maintenance) and per-operating analgesia (Before surgery: Buprenorphine 0.05mg/kg IM 30 minutes, local cutaneous injection of lidocaine around the abdominal aperture; after surgery: renewal of buprenorphine 4 hours later, then meloxicam PO for 2 days). At the end of the experiment, the implants were removed by the same procedure. This method allowed to record individuals’ body temperature (Tb) every 5 minutes. Tb were analyzed as minimal Tb registered each day (‘Tbmin’, in °C), and were computed as a mean during the CTL and caloric treatment phases of the experiment. We also measured the duration of Tb under 33°C (torpor duration, ‘TD’ in minutes), a threshold often used in torpor analysis (Giroud et al., 2008; Canale et al., 2011).

#### Markers of physiological stress

Cortisol (ng.ml^− 1^; Cortisol Urine ELISA from LDN®, ref MS E-5100) and 8-hydroxy-2’-deoxyguanosine (“8-OHdG,” in ng.ml^− 1^; OxiSelectTM Oxidative DNA Damage Elisa kit, Cell Biolabs Inc., ref STA-320) were measured in duplicates from urine samples as indicators of the organisms’ response to environmental stress (Miller et al., 1991) and oxidative-stress related DNA damage (Loft et al., 1993), respectively. Creatinine concentration (mg.ml^− 1^) was used to normalize all urine measurements as an indicator of renal filtration activity (MicrovueTM Creatinine Elisa kit, Quidel® Corporation, ref 8009). Results are thus expressed in ng.mg Creat.^− 1^.

#### Statistical analysis

Results shown are given as medians ± standard deviation (s.d). No outlier was identified or removed from data sets after testing with Dixon’s Q test. Statistical analysis was conducted by the use of R software v 3.5.1 (R Core Team, 2016), and tests were considered significant when p-values were below the significant level set at 0.05. We applied linear mixed models with random individual effect to test the effect of sex, the phase of the experiment (before and after two weeks of caloric treatment), caloric treatment (CTL vs. CR) and their interaction. Body mass was included in models only when its effect was significant on the explained variable. Non-parametric pairwise Wilcoxon test was used as post-hoc analysis (pairwise nonparametric tests for caloric treatment effect on the same individuals, unpaired for sex effect or tests on variations in %). Medians and standard deviations are presented along with statistical parameters. The principal component analysis was performed on the variations of each parameter (%) after caloric treatment (either CTL or CR rations). We used the “FactominR” 1.34 package (Lê et al., 2008), and missing values were imputed with the “missMDA” 1.11 package (Josse and Husson, 2016).

## 3. Results

### Sex-specific effect of acute CR on BM in late winter

At the beginning of the experiment, BM did not statistically differ between males and females (Females= 129 ± 11.5 g; Males= 110 ± 13.9 g; W=82, p=0.17), and even when BM was reported to body length (Females= 9.0 ± 0.8 g.cm^-1^; Males= 8.5 ± 1.2 g.cm^-1^; W= 53, p=0.85), considering that males are shorter than females (Females= 14.3 ± 0.3 cm; Males= 13.5 ± 0.3 cm; W=92.5, p=0.001). The effect of CR treatment on BM significantly depended on sex (Sex:Trt:Phase: Chisq=8.21, p= 0.0042; Figure 1A). Furthermore, post-hoc analysis showed that even though CR females lost about 14% of their initial BM, only the 21 % BM loss of CR males was significant (CTL Females = + 1.5 ± 5.5 %, W= 1.5, p= 0.14; CR Females= -14.5 ± 3.4 %, W=15, p=0.057; CTL Males = + 9.6 ± 5.6 % W=1, p= 0.13; CR Males = -21.5 ± 2.8 %, W= 21, p=0.036). The BM loss between CR males and CR females was also significantly different (W=30, p=0.0043).

**Figure 1.**
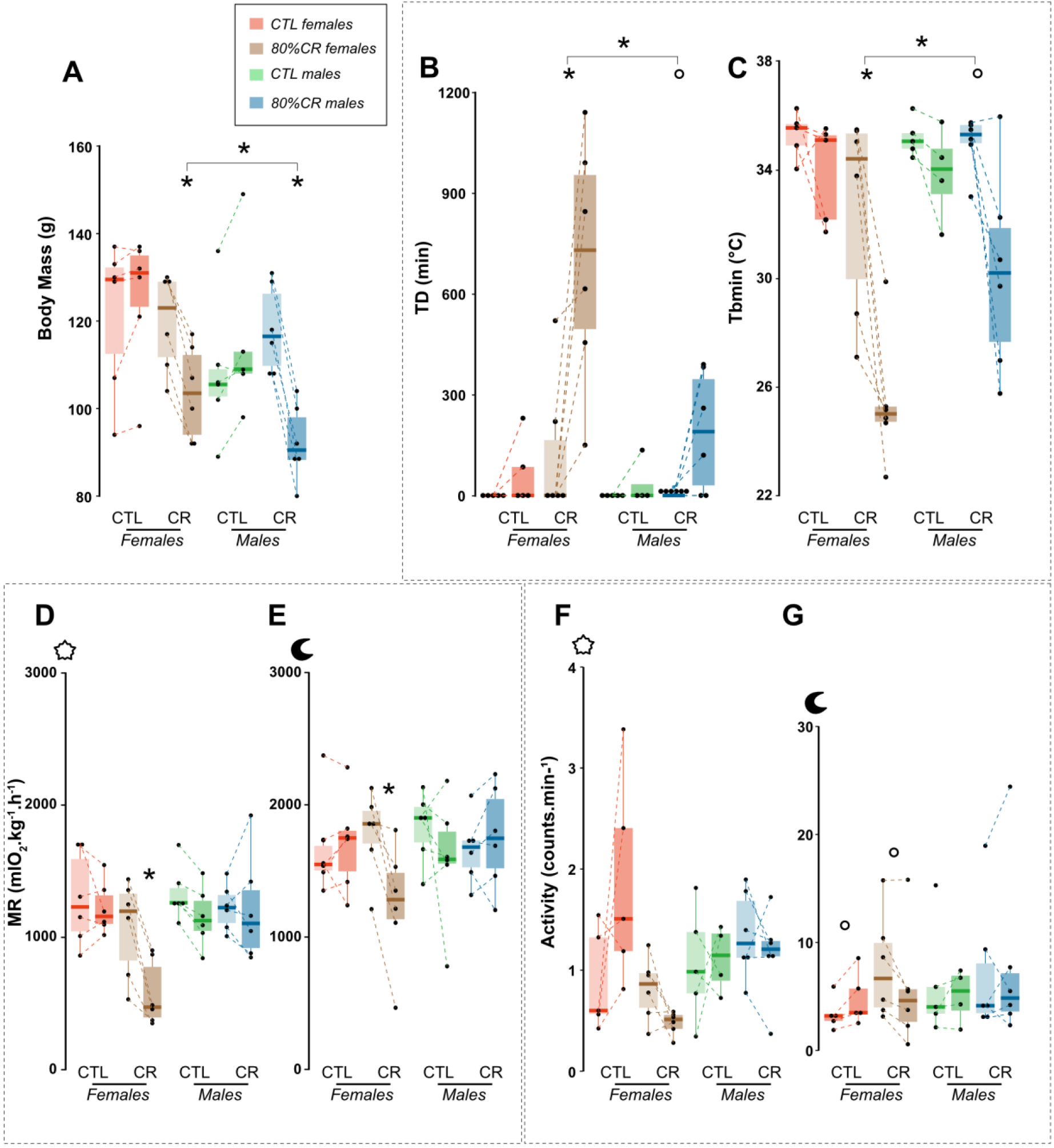
Overview of the thrifty female. **A:** Body mass (g) in each group (CTL females in orange, CR females in brown, CTL males in red, CR males in blue) before (lighter colour) and after (darker colour) caloric treatment (either CTL treatment or 80% CR). B: Torpor duration (minutes); C: daily minimal body temperature (Tbmin, in °C). D: mean MR (mlO_2_.kg^-1^.h^-1^) during the day. E: mean MR (mlO_2_.kg^-1^.h^-1^) during the night. F: Mean activity (counts.min^-1^) during the day. G: Mean activity during the night. Intra-group effect are analysed with a non-parametric pairwise Wilcoxon-Mann-Whitney test, and differences between CR groups are come from linear mixed models. *<0.05; °<0.1.

### Sex-specific torpor response to CR

As for BM, males and females showed different increase in torpor duration in response to CR (Sex:Trt:Phase: Chisq= 3.1, p=0.08, Figure 1B). Indeed, torpor duration significantly increased in CR females (+650 ± 397 minutes, V=0, p= 0.031), and at a lesser extent in CR males (+178 ± 179 minutes, V=0, p= 0.1). Minimal Body temperature also decreased in all CR animals, but at a lesser extent in males (CR females = -22.2 ± 10.5 %, V=21, p=0.03; CR males = -12.0 ± 10.6 %, V=20, p= 0.06; Figure 1C). Considering data distribution (non-gaussian, negatively skewed), this effect could not be confirmed in a linear mixed model (Sex:Trt:Phase: Chisq= 0.75, p= 0.39).

CR had a sex-specific effect on MR during night and day (Sex:Trt:Phase during day: Chisq= 15.7, p < 0.0001; during night: Chisq= 11.2, p= 0.0008). MR was only decreased after CR in females during the day (CR females = -44.4 ± 13.4 %, V=21, p= 0.004) and during the night (CR females = -34.9 ± 20.0 %, V=21, p= 0.03); while no change in MR levels were observed in CR males during both periods (CR males = -8.4 ± 20.8 %, V=12, p=0.84 during the day; +9.0 ± 14.2 %, V= 6, p= 0.44 during the night).

### Sex-specific activity patterns in response to acute CR in late winter

Mean activity during the day tended to differ between sexes depending on treatment (Sex:Trt:Phase: Chisq= 2.78, p= 0.096; Figure 1F), and this effect of CR was more marked in females (CTL Females = +92 ± 210 %, V= 2, p=0.19; CR females = -42 ± 39 %, V=18, p = 0.16), than in males (CTL males = +29 ± 150 %, V= 3, p=0.63; CR males = -18 ± 37 %; V = 15, p= 0.44). During the night, mean activity tended to differ between sexes depending on treatment (Sex:Trt:Phase: Chisq = 3.5, p = 0.06; Figure 1G) with a significant decrease of activity only in CR females (CTL females = +38 ± 26 %, V=0, p=0.06; CR females = -36.3 ± 27.5, V = 20, p = 0.06; CTL males = +10.3 ± 58.8 %, V=1, p=0.25, CR males = +21.3 ± 49.3 %, V=8, p= 0.69).

### Principal Component Analysis on thrifty female’s characteristics

The PCA based on the changes observed after the caloric treatment showed significant discrimination of CR females along the first axis explaining 55.7% of the data (Figure 2A and 2B). CR females discriminated from CTL females with greater change in torpor bouts duration and depth (lower Tbmin). They were also characterized by lower variations in day and night activity, MR, and BM. In contrast, CR males were discriminated from the others on the basis of a greater BM loss (in a stronger way than for CR females, see figure 2B) and a lower change in activity during the day.

**Figure 2.**
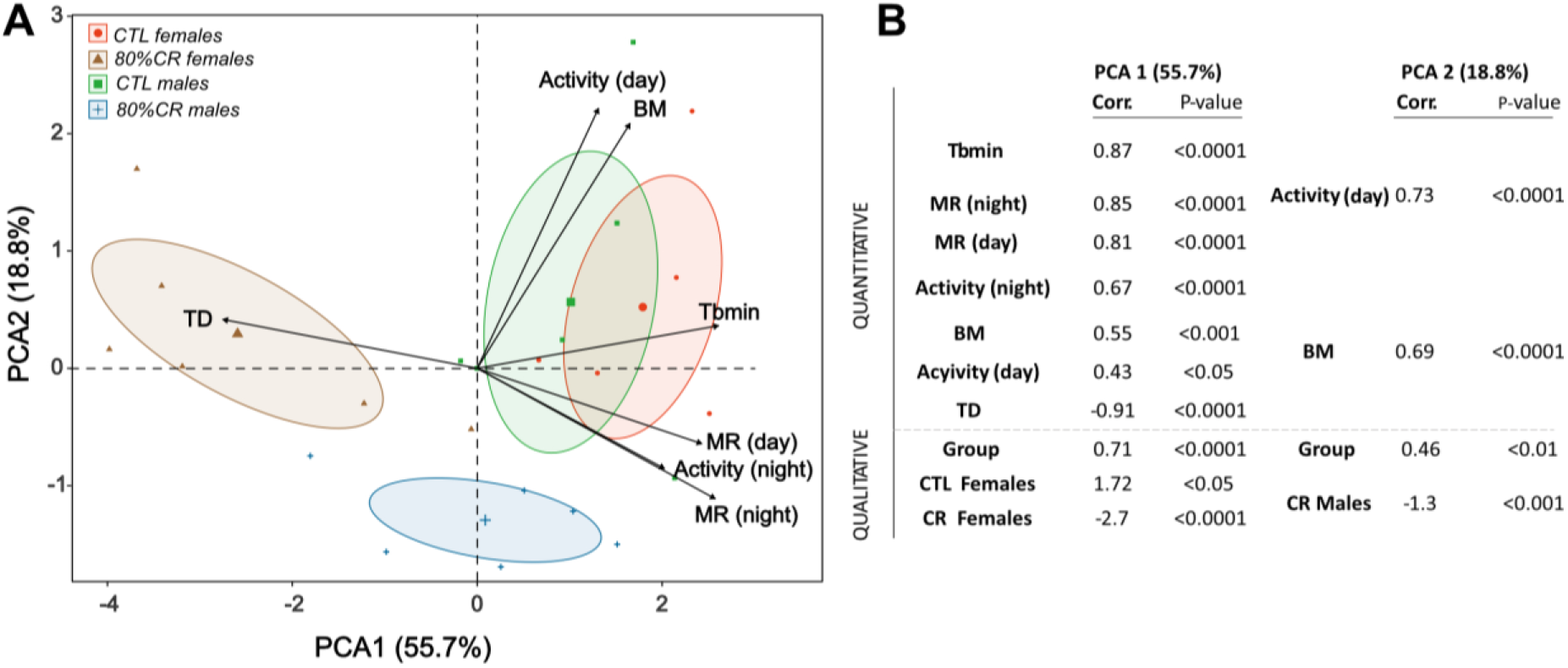
Principal Component analysis discriminating male and female mouse lemurs submitted or not to CR, by the variation of their parameters (in %) before and after caloric treatment (CTL, green and orange for males and females, respectively; CR, blue and brown for males and females, respectively). Parameters indicative of energy balance (Body mass “BM”), metabolic rate (Mean MR over day “MR (day)”, Mean MR overnight “MR (night)”, Torpor duration “TD”, Daily minimal body temperature ‘Tbmin”), and activity pattern (Mean activity during day “Activity (day)”, and Mean activity during night “Activity (night)”) were included in the analysis. Individual plots represent the variation of each parameter in response to the caloric treatment, in %. They are shown in (A) and grouped into sex*Trt. The contribution of each parameter is also represented by black arrows displaying the “Variable factor map.” Principal Component 1 (PC1) indicates ∼56% of variability and Principal Component 2 (PC2) ∼19%. (B) Table of correlation coefficients and corresponding p-values shows only the variables significantly contributing to the two first principal components of the analysis (Quantitative) and the discrimination of qualitative variables: CTL Females, 80%CR Females, CTL Males, 80%CR Males.

### Cortisol and oxidative damage evolution after CR

The decrease in cortisol levels after CR did not appear significant in both males and females in the linear mixed model (Figure 3A; Phase:Trt: Chisq= 1.49, p= 0.22; CR females= -6.6 ± 58.4 ng. mgCreat^-1^, V=9, p=0.81; CR males= -24.0 ± 92.5 ng. mgCreat^-1^, V=13, p=0.69). Oxidative damage did significantly decrease with caloric treatment however, independently of the sex (Phase:Trt: Chisq= 5.47, p=0.019; Phase:Trt:Sex: Chisq= 0.0046, p=0.83; Figure 3B). Additionally, initial 8-OHdG concentrations were overall higher in males (Females= 29.04± 10.9 ng. mgCreat^-1^, Males= 37.7 ± 16.8 ng. mgCreat^-1^; Sex effect: W=130, p=0.014; Figure 3B), and the levels after CR were still higher in CR males as compared to CR females (CR females =25.2 ± 3.3 ng. mgCreat^-1^; CR Males= 33.1±7.9 ng. mgCreat^-1^, W=3, p=0.03).

**Figure 3.**
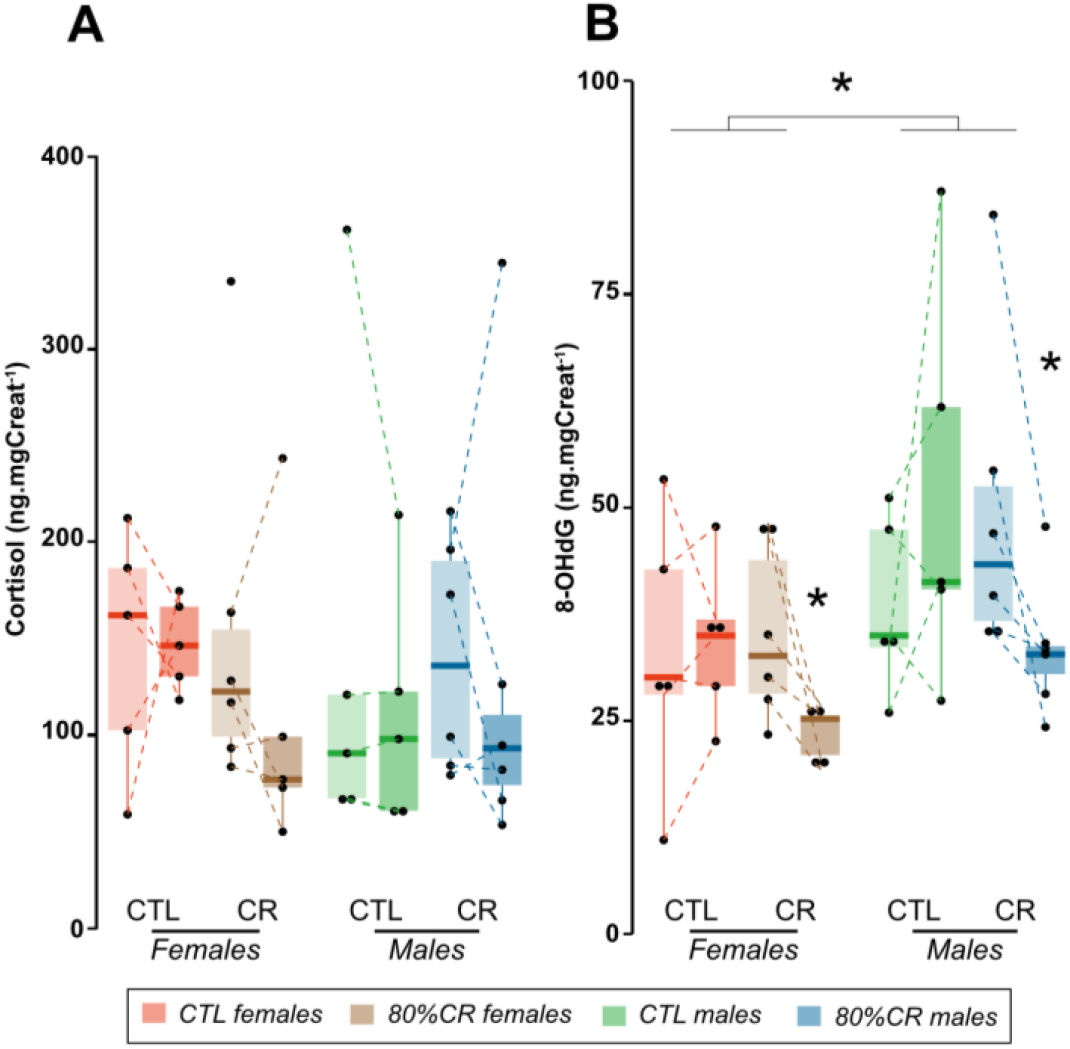
Urinary concentrations of A: Cortisol (ng.mgCreat^-1^); B: 8-OHdG (ng.mgCreat^-1^); in each experimental group (CTL females in orange, CR females in brown, CTL males in red, CR males in blue) before and after caloric treatment (either CTL treatment or 80% CR).

### Effect of CR on the reproductive axis

CR females did not delay their estrus after entering LD photoperiodic regimen, as each CTL and CR groups showed estrus around the 17e day after LD transition (CTL females = 18 ± 3.8 days, CR females= 17 ± 1.9 days, W=11, p=0.9). In males, testis grew in size between the first and the second phase of the experiment, but CR had no effect on it (+ 50 ± 0.35 % for both CTL and CR males, W=18.5, p=0.57).

## 4. Discussion

The thrifty female hypothesis (TFH) is associated with seasonal reproduction, which shapes intersexual differences in energy balance. In its first description in wintering little brown bats, the TFH characterised lower BM loss in wintering females as compared to males (Jonasson and Willis, 2011). Here we explored thrifty phenotypes in *Microcebus murinus*, by looking at differences in BM loss after CR, sex specificities in body temperature modulations and torpor use, in relation with metabolism and activity patterns.

### *Confirming the Thrifty Female Theory in* Microcebus murinus

After an intense 80% caloric restrictions, female grey mouse lemurs lost significantly less BM than males, which confirms a sex gap in energy balance. Moreover, we confirmed the link between BM sex-specific modulation and substantial diminution in body temperatures by showing that females used significantly longer and deeper torpor bouts than the males under CR and the other animals under a CTL diet. Torpor is defined by low body temperature, which follows decreasing basal metabolic rate (BMR) in a relationship characterized by hysteresis (Geiser et al., 2014). Hypometabolism is the energy saving process used by heterotherms, either daily or over the season, that limits energy expenses in a challenging context of low food availability or low ambient temperatures that would deeply impact caloric loss. Here only CR females showed significant diminution in MR (VO_2_) after treatment, correlated with Tb modulations and diminution in day and night activity (see ACP Figure 2). In consequence, we proved that females adapt their metabolism and behaviour to food shortage on different levels, and that this modulation has a relative positive impact on their energy balance as compared to males which did not show diminution in MR and lost more BM. Here, we thus confirmed that female *Microcebus murinus* match the TFH, as it is defined in *Myotis lucifugus* (Jonasson et Willis 2011). As opposed to females, even if males showed a substantial -yet inferior-decrease in body temperatures after CR, this was not associated to a decrease in metabolic rate. Hence, males may present additional costs in using torpor than females, or less efficacy in saving energy by using it.

The seasonal reproductive cycle of the little brown bat is however very different from the one of grey mouse lemurs. Indeed, these chiropteras’ mating period takes place at the end of summer and females delay ovulation and fecundation by storing sperm in sacs throughout winter (Fenton and Barclay, 1980; Wai-Ping and Fenton, 1988). This “non selective mating” process allows males to mate with “passive” hibernating females during winter (Wai-Ping and Fenton, 1988), a fact that underlines very clearly the gap between sexes in metabolic activity during the poor season. Moreover, bats use heterothermy to survive winter, especially since they have the particularity to display large wings responsible for high levels of heat loss (Dzal and Brigham, 2013). However, torpor has costs, from dehydration to memory loss, increased vulnerability to predators and temporary diminished immune function (Luis and Hudson, 2006; Bieber et al., 2014; Wei et al., 2018; Landes, 2020). These facts led to the “torpor optimization hypothesis”, which states that torpor should be used only if fat stores are required for survival and/or reproduction; it would thus be regulated in its intensity according to the energy allocation required (Humphries et al., 2003). The hypothesis (and verified fact) that females little brown bats are thriftier than males during winter is based on this trade-off between reproduction optimization and survival costs associated with torpor. Based on Jonasson and Willis (2011), predictions are that adult females maximize energy saving for reproduction by using deeper torpor, while males aim at avoiding the physiological or ecological secondary costs of torpor by reducing its intensity and therefore and using fat reserves more quickly.

In contrast to little brown bats, grey mouse lemurs mating takes place during the transition to the wet season in nature (Radespiel et al., 2006), and about 2 weeks after LD initiation in laboratory conditions (Perret and Aujard, 2001). Even if males *Microcebus murinus* begin spermatogenesis earlier during the second half of winter to prepare for mating (Perret et Aujard 2001, Noiret et al. 2021), the expected gap of MR between sexes is not so obvious compared to the little brown bats, as females stay active during the night. Female birds and mammals are known to reactivate their reproductive axis by responding to photoperiodic change when transitioning to summer (Simonneaux and Ribelayga, 2003). In females *Microcebus murinus*, even if they synchronize with photoperiodic transition to LD to fully reactivate their reproductive activity, they do maintain an endogenous circannual sexual cycle as proven by the expression of oestrus around 30 weeks after constant exposition to SD in laboratory conditions (Perret and Aujard, 2001; Hozer and Perret, 2023). However, we do not know if they anticipate the reproductive season by reactivating their metabolism during late winter as males do, which would tighten the gap in energy expenditure between the two wintering sexes. Indeed, a previous follow up of the effects of a chronic (6 months) and moderate (40%) CR applied in wintering grey mouse lemurs showed that females lost as much BM as CR males during the second half of winter contrary to what was expected, and despite different torpor profiles in response to CR (Noiret et al, 2021). The present study however shows that females *Microcebus murinus* exhibit thriftier phenotypes than males under more acute and intense conditions during late winter, which conforms with the TFH.

Contrary to Johansson and Willis however, and based on our observations in *Microcebus murinus*, we think that males do optimize reproductive success in using less torpor than females. We know that spermatogenesis implies shorter torpor bout in order to maintain optimum metabolism and temperature to promote gamete production (Barnes et al., 1986; Gagnon et al., 2020), especially since males *M. murinus* display features associated with sperm competition (Aslam et al., 2002; Simmons, 2005). In consequence, we think that females’ thrifty phenotype either could be a result of a contrast to males’ greater allocation to reproduction during winter (and the double cost of higher metabolic rate in a context of low food availability) with a subsequent greater BM loss, or else could derive from better fitted adaptive response in females. Indeed, as Jonasson and Willis state, stronger selective pressures would act on females’ reproduction, leading to a sex-specific adaptation in energy saving mechanisms that would benefit females in the trade-off between short-term survival (i.e. torpor costs) and reproduction. As a matter of fact, both mechanisms -negative energy balance in males, and better energy sparing capacities in females-could intervene in shaping the female thrifty phenotype. Hence, if seasonality selected for better energy saving mechanisms in females, it would mean that sexes are unequal regarding the classic reproduction-survival trade-off. Because torpor has costs, Jonasson and Willis based their sex-variability theory on the idea that it should be avoided as much as possible (Humphries et al., 2003). Our results suggest that females would host better active machinery specialized in reducing the costs of torpor, as it was shown in summer (Noiret et al., 2020). Yet, torpor has costs but also benefits (Geiser, 1998, 2004; Keil et al., 2015), especially for surviving winter. Under the TFH, seasonally breeding females would thus advantage both reproduction and survival by using torpor, while males would ultimately impact survival as they have to “wake up” to invest in reproduction.

Also, because we strongly link the TFH with seasonal breeding, one may oppose the observation of thriftier females in non-seasonal breeders, generally domesticated species that lost photoperiodicity in a seasonally buffered environment due to human activity (Setchell, 1992; Blottner and Jewgenow, 2007). Indeed, female laboratory rats showed greater thermogenic functionality of their brown adipose tissue compared to males, and a higher resistance to weight loss (Rodríguez-Cuenca et al., 2002; Valle et al., 2007), features that highly resembles thrifty phenotypes, and are shared with female seasonal breeders (Lennox and Goodship, 2008; Terrien et al., 2010; Kart Gür and Gür, 2015). As males and females rats endure rapid and continuous spermatogenic and oestrous cycles respectively throughout the year (around 50 days for a male and 4 to 5 days in females with no seasonal discontinuity; (Lohmiller et al., 2006), we believe that the thrifty phenotype is a sex-specific adaptation to delayed reproductive allocation and big energy expenses in females, that would be an inherited residue -still beneficial-in non-seasonal breeders.

### On sex-specific impact of “unexpected” CR in late winter and during reproductive investment

It is somehow surprising, considering the present results, that a chronic 40% CR could not discriminate a thriftier phenotype in female grey mouse lemurs (Noiret et al, 2021). In comparison, this 80% intense CR was designed to be considered as an acute stress compared to the CTL ration animals received during their upstream winter in breeding conditions, before the start of the present experiment. This fact could be the factor triggering male and female differences in energy balance. However, an acute winter CR would not necessarily be a stressor in the context of the highly seasonal and hypervariable environment like Madagascar, to which our animals are adapted (Dewar and Richard, 2007). In addition, this 80% CR did not result in cortisol elevation, as compared to the previous 60% CR we applied in post-reproductive LD acclimated animals (Noiret et al., 2020), were all CR individuals showed high cortisol increase after food shortage. Nonetheless, both situations led to a decrease in oxidative damage in urine when corrected with urinary creatinine and males presented higher levels of oxidative damage than females, as it was already observed in the 60% CR study (Noiret et al., 2020). Added to their lower body mass in late winter, and considering the high energy expenses they will face during spring in the wild, male *Microcebus murinus* survival chances may be reduced compared to females. In nature, males’ survival at the onset of summer was shown to be 16% inferior to the one of females, but this effect is usually attributed to more “risky” behaviours in males (Kraus et al., 2008), with territorial defence, male-to-male competition and the exposition to predators that comes with it. Even if recent observations of wild populations do not show any sex-bias, we cannot exclude the phenomenon in the future, as more and more unpredictable environmental events are expected to occur with global warming, that could challenge males more than females (Canale et Henry 2010; Walter et al. 2011; Easterling 2000). It has been pointed out that even if sexual reproduction is at the core of many ecological and evolutive theories, sex has been overlooked in population dynamics (Bessa-Gomes et al., 2010), and two-sex models are in fact more accurate to predict extinction risk compared to the usual one sex-models (Caswell and Weeks, 1986). In regard of the catastrophic consequences a sex-bias in mortality could have for population extinctions (Wedekind, 2002; Galliard et al., 2005; Grayson et al., 2014), it would be a matter of importance to assess more acutely sex differences in energy-saving strategies and its impact on fitness, to better predict the future of wild population and act in consequence.

### Conclusion

We provided evidence for the TFH in female *Microcebus murinus* after intense CR in late winter, at the onset of the reproductive period. This phenomenon is probably linked to a sex-gap in energy allocation to reproduction due to seasonality, males investing earlier than females, as well as a sex-specific adaptation in energy saving mechanisms linked to greater selective pressures concerning reproductive success in females. After being proved in bats and primates, the TFH seems quite robust. In an evolutive point of view, the TFH would mean that by using torpor, females would promote both reproduction and survival in a seasonal context, while males have to endure a more classic unbalanced trade-off: as they invest in reproduction, they ultimately disadvantage survival. Males showing poorer energy balance before mating, especially in a context of climate change, could face bigger loss than females, leading to sex-biased population with an increased extinction risk.

